# Disuse-driven plasticity in the human thalamus and putamen

**DOI:** 10.1101/2023.11.07.566031

**Authors:** Roselyne J. Chauvin, Dillan J. Newbold, Ashley N. Nielsen, Ryland L. Miller, Samuel R. Krimmel, Athanasia Metoki, Anxu Wang, Andrew N. Van, David F. Montez, Scott Marek, Vahdeta Suljic, Noah J. Baden, Nadeshka Ramirez-Perez, Kristen M. Scheidter, Julia S. Monk, Forrest I. Whiting, Babatunde Adeyemo, Abraham Z. Snyder, Benjamin P. Kay, Marcus E. Raichle, Timothy O. Laumann, Evan M. Gordon, Nico U.F. Dosenbach

**Affiliations:** Department of Neurology, Washington University School of Medicine, St. Louis, MO 63110; Department of Neurology, New York University Grossman School of Medicine, New York, New York 10016, USA; Basque Center on Cognition, Brain and Language, Donostia, Gipuzkoa, Spain; Department of Biomedical Engineering, Washington University in St. Louis, MO 63130; Division of Computation and Data Science, Washington University School of Medicine, St. Louis, MO 63110; Department of Psychiatry, Washington University School of Medicine, St. Louis, MO 63110; Mallinckrodt Institute of Radiology, Washington University School of Medicine, St. Louis, MO 63110; Department of Psychological and Brain Sciences, Washington University in St. Louis, St Louis, MO, USA; Department of Neuroscience, Washington University School of Medicine, St Louis, MO, USA; Department of Pediatrics, Washington University School of Medicine, St. Louis, MO 63110

## Abstract

Motor adaptation in cortico-striato-thalamo-cortical loops has been studied mainly in animals using invasive electrophysiology. Here, we leverage functional neuroimaging in humans to study motor circuit plasticity in the human subcortex. We employed an experimental paradigm that combined two weeks of upper-extremity immobilization with daily resting-state and motor task fMRI before, during, and after the casting period. We previously showed that limb disuse leads to decreased functional connectivity (FC) of the contralateral somatomotor cortex (SM1) with the ipsilateral somatomotor cortex, increased FC with the cingulo-opercular network (CON) as well as the emergence of high amplitude, fMRI signal pulses localized in the contralateral SM1, supplementary motor area and the cerebellum. From our prior observations, it remains unclear whether the disuse plasticity affects the thalamus and striatum. We extended our analysis to include these subcortical regions and found that both exhibit strengthened cortical FC and spontaneous fMRI signal pulses induced by limb disuse. The dorsal posterior putamen and the central thalamus, mainly CM, VLP and VIM nuclei, showed disuse pulses and FC changes that lined up with fmri task activations from the Human connectome project motor system localizer, acquired before casting for each participant. Our findings provide a novel understanding of the role of the cortico-striato-thalamo-cortical loops in human motor plasticity and a potential link with the physiology of sleep regulation. Additionally, similarities with FC observation from Parkinson Disease (PD) questions a pathophysiological link with limb disuse.

## Introduction

Brain networks must simultaneously exhibit stability, to preserve acquired skills, but also flexibility, to learn and adapt to environmental changes^1^. Motor behavior is generated by complex cortico-subcortical circuits including the thalamus, basal ganglia and cerebellum^2–4^. Cortico-striato-thalamo-cortical loops have been intensively studied to understand pathways involved in motor learning. Prior studies of plasticity in this circuit have been pursued largely by invasive electrophysiology in patients undergoing deep brain stimulation [DBS])^5–7^, and animal models^8,9^. Human neuroimaging studies of motor plasticity have focused on the cerebral cortex because of the low signal-to-noise ratio of blood oxygen dependent (BOLD) signals in subcortical structures ^10,11^ and the limited effect sizes typically seen in plasticity paradigms^12–14^.

To study plasticity mechanisms in humans, we developed an experimental paradigm that induces disuse by restraining the upper-extremity using a full length cast ^15–17^. This approach is similar to classical animal plasticity studies, which impose motor or sensory restrictions (e.g., limb constraint, deafferentation, monocular deprivation) in a small number of intensively studied individuals ^18,19^. We casted the dominant (right) upper extremity of three participants (Nico, Ashley and Omar) for two weeks and collected around-the-clock actigraphy and daily task and resting state functional MRI (fMRI) over a 6-week experimental protocol (2 weeks pre-cast, 2 weeks casting, 2 weeks post-cast). Limb disuse was documented by the actigraphy data and showed 15 to 24% increased use of non-dominant hand use. The participants only exhibited a reduced grip strength of the casted extremity measured at cast removal and recovered within days with no persistent deficits.

Dense longitudinal fMRI sampling enabled us to perform within participant analyses using our individual-specific Precision Functional Mapping (PFM) methodology ^20^. Resting state functional connectivity (FC) noninvasively maps functional networks within the brain and how they change in response to disuse. Previous analyses of the fMRI data acquired in the casting experiment focused on the cerebral cortex ^16,17^. FMRI signals in the left and right upper extremity primary somatomotor cortex (SM1_ue_) are strongly correlated at baseline. Casting induced a marked decrease in FC (-0.23 to - 0.86 change in correlation from around 0.8 before casting) restricted to the upper extremity specific parts of SM1_ue_. We also detected increased FC between left SM1_ue_ and the cingulo-opercular network (CON)^16^, an executive control system responsible for initiating and maintaining actions ^21,22^. Unexpectedly, high amplitude fMRI signal pulses arising in left SM1_ue_ emerged during the casting period and were detected in supplementary motor area and cerebellum ^17^.

Recently, we also described the previously unrecognized Somato-Cognitive Action network (SCAN), which is interleaved with effector-specific motor regions along the central sulcus ^23^. The SCAN is strongly functionally connected to the CON, and seems to serve as its downstream actuator, turning abstract plans into integrated whole-body actions. Becoming aware of the sharp divisions between SCAN and effector-specific motor regions also provided motivation to re-evaluate our prior interpretations of some of the upper-extremity disuse-driven plasticity effects.

Motor and action control cannot be understood without subcortical nodes. Basal ganglia (putamen, caudate, globus pallidus, etc.) and thalamic nodes are intricately networked to facilitate the selection of intended actions while simultaneously suppressing or inhibiting potentially conflicting or undesirable ones ^24^. The posterior putamen specifically has been shown to be important for slow but long-term establishment of habits ^25,26^.

The central thalamus also plays a significant role in motor adaptation and control by serving as a relay and integration center between various brain regions involved in motor functions ^27^. The thalamus comprises a multitude of distinct nuclei, including the ventralis intermedius (VIM), the centromedian (CM), and the ventroposterior lateral (VPL) ^28,29^. The VPL receives sensory information and relay information for fine tuning of movement^29^. The VIM, in comparison, is primarily involved in motor control functions such as planning, initiation, and execution of voluntary movements^30^. The VIM is the deep brain stimulation (DBS) target in Essential Tremor and tremor-predominant Parkinson’s Disease (PD) ^31–35^. A case report from a patient with longstanding bilateral upper extremity loss undergoing DBS suggested plasticity of VIM neurons, leading to an over-representation of shoulder movements^36^.

The centromedian nucleus (CM) of the thalamus has been classified as part of the ‘non-specific’ nuclei of the thalamus and shows specific projections to the sensorimotor regions and anterior cingulate ^8^. The CM plays an important role in the regulation of the cortical parvalbumin neurons that promote Hebbian plasticity ^37–40^, and the overall level of cortical activity and arousal ^41^. CM has mainly been targeted with DBS in refractory generalized epilepsy ^42–44^ due to its role in sleep-wake regulation and arousal. The central thalamus plays an important role in regulating sleep stages and generates sleep spindles important for memory consolidation ^45–47^, including procedural memory.

Given the importance of subcortical structures for motor control and skill learning, we expanded our prior analyses to investigate disuse-driven plasticity in human subcortical circuits for the first time.

## Results

### Disuse strengthens motor cortex functional connectivity with central thalamus and posterior putamen

Measures of FC in the subcortex are smaller than in the cerebral cortex. This observation is, at least partly because fMRI signal-to-noise ratio drops off towards the center of the brain as distance from the MRI coil elements increases. Consequently, the magnitude of FC changes between different conditions (e.g., Cast - Pre) are greater in the cortex than in the subcortex. Therefore, to measure FC changes in the subcortex, we used standardized effect size (Cohen’s d) to account for the signal-to-noise differences. The present effects, expressed in terms of Cohen’s d, match our previously reported parcel-based FC findings^16^. Use-driven FC changes (Cohen’s d) of the left SM1_ue_ increases with the CON regions (purple border, figure S1) for all three participants in cortex and with hand motor regions in the cerebellum (Figure S1, S3). We note that disuse-driven FC changes spared the Somato-Cognitive Action Network (SCAN; Figure S1, maroon outlines), a set of effector-general action control regions in motor cortex^23^ strongly functionally connected with the CON (see also Figure S2). Thus, the two SCAN regions around the upper-extremity specific primary motor region (black seed region, left SM1_ue_) did not show significant FC changes due to casting.

In the subcortex, all three participants showed statistically significant increases in FC between disused left SM1_ue_ and the central thalamus, as well as the posterior putamen (Figure 1, bottom row; cluster-based thresholding; see Methods). The subcortical effects (Cohen’s d) were comparable in magnitude to effects observed in SM1_ue_ (Table 1).

**Figure 1:**
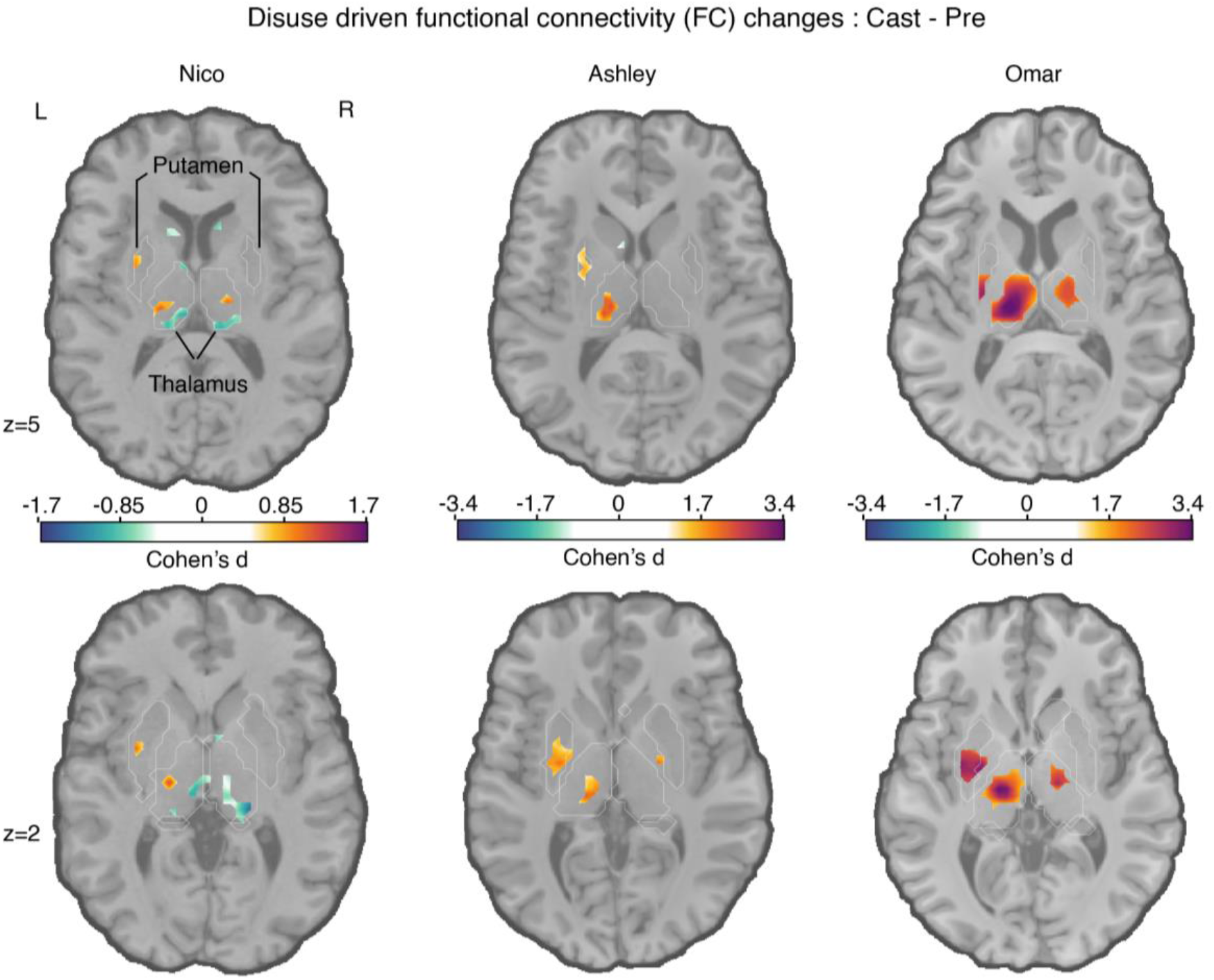
Disuse-driven changes in functional connectivity (FC) of the left effector-specific primary somatomotor cortex (L-SM1_ue_) in subcortex. Individual-specific plasticity effect size (Cohen’s d) maps showing changes in FC during right arm casting (Cast − Pre) for the L-SM1_ue_ for each participant (left to right columns: Nico, Ashley, Omar). For reference, a Cohen’s d of 0.8 is generally considered a large effect size. Only significant effects after cluster correction at p < 0.05 (see Methods) are displayed. Please note, Nico’s data were collected using an earlier pulse sequence with a TR that was twice as long (2.2 s) compared to Ashley and Omar (1.1s). Nico’s effect sizes are about half the size of the other participants. The FreeSurfer based anatomical borders of the putamen and thalamus are shown as white outlines (bottom row).

**Table 1:** Disuse-driven functional connectivity changes in subcortex. Peak coordinate (x,y,z) in MNI space and cluster size (N_vox_) for subcortical regions with disuse-driven increases in functional connectivity with L-SM1_ue_. as well as maximum Cohen’s d values (Max) of the left putamen, left thalamus, cortex and cerebellum

|  | Nico | Ashley | Omar |
| --- | --- | --- | --- |
| Left posterior Putamen | -31 -6 0<br>N<sub>vox</sub> = 43<br>Max = 1.17 | -25 -12 0<br>N<sub>vox</sub> = 76<br>Max = 2.56 | -31 -9 3<br>N<sub>vox</sub> = 71<br>Max = 3.09 |
| Left Thalamus | -16 -21 6<br>N<sub>vox</sub> = 68<br>Max = 1.24 | -7 -21 0<br>N<sub>vox</sub> = 72<br>Max = 2.83 | -13 -24 6<br>N<sub>vox</sub> = 219<br>Max = 3.86 |
| Cerebellum | Max = 2.22 | Max = 2.68 | Max = 4.79 |
| Cortex | Max = 2.43 | Max = 4.01 | Max = 4.49 |

Other FC changes were significant in some but not all participants. In particular, in Ashley and Omar, a small area of significant FC increase was observed in the left posterior globus pallidus bilaterally. Ashley and Omar’s effect sizes were roughly twice those observed in Nico. These differences could be attributable to inter-individiual differences, or due to differences in the fMRI pulse sequence (TR 1.1s Ashley, Omar vs. 2.2s Nico), or fMRI signal signal-to-noise ratios ^48^.

### Disuse pulses in the central thalamus and motor cerebellum

We previously reported the emergence of large fMRI signal pulses in the disused motor cortex after 12-48 hours of arm casting. To identify disuse pulses in the subcortex, we developed an HRF-based pulse detection method that accounts for spatial variability in HRF shape (see Methods). Using the HRF-based detection method, we were able to detect the presence of disuse pulses in the subcortex on top of confirming their spatial distribution in the cortex (Figure 2; left) and cerebellum (Figure 2; third column) ^17^. Disuse pulses in the central thalamus were observed in all participants (Figure 2; second column). The peak pulse percent signal change in the thalamus was lower compared to cortex (Figure 2, right column). In the participant with the most disuse pulses (Ashley), we also detected them in the posterior putamen. The least number of pulses were detected in Nico, who also exhibited the smallest FC changes.

**Figure 2:**
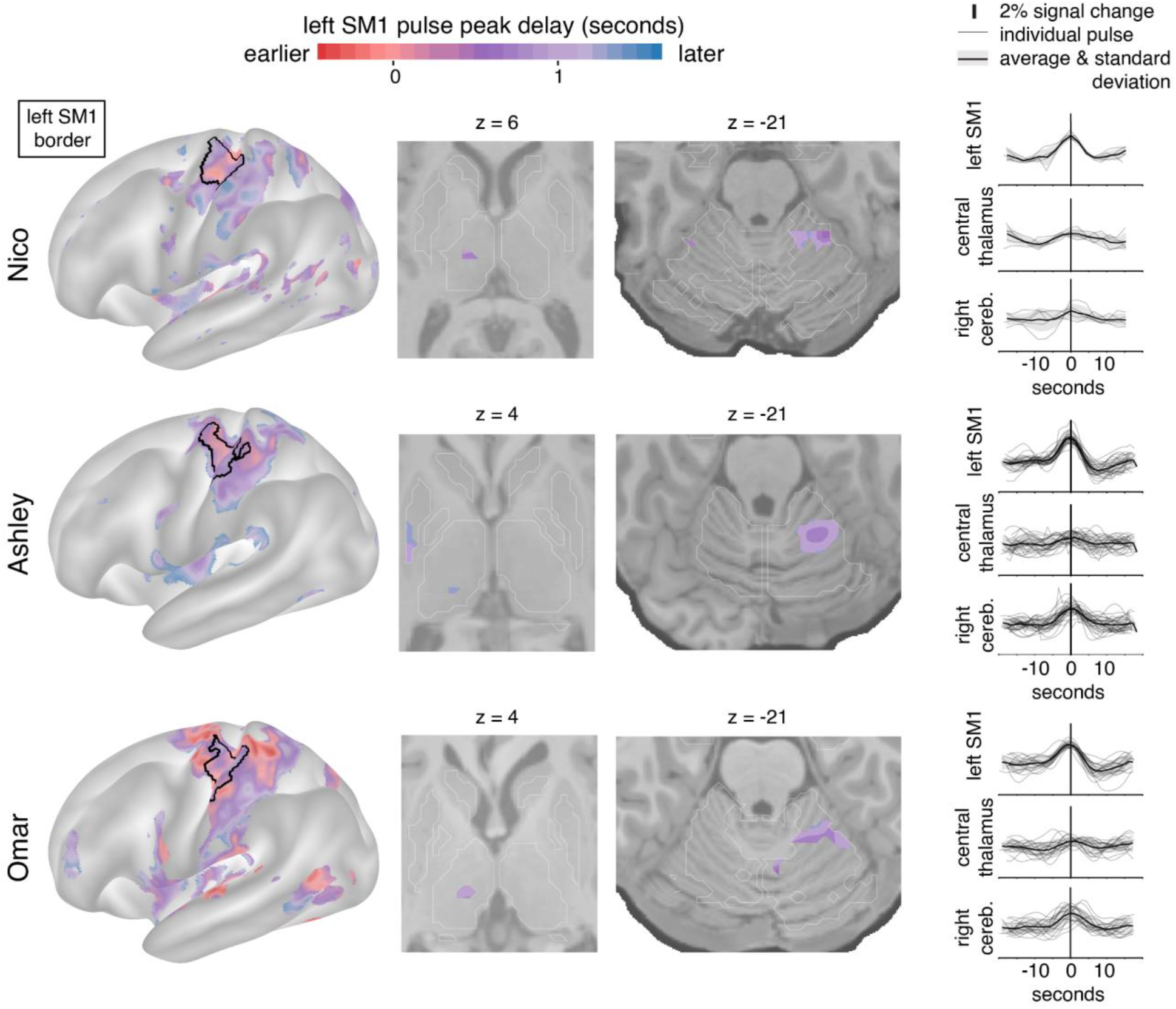
Disuse pulse distribution in cortex and subcortex. The timecourse of each disuse pulse observed during casting was modeled using a Hemodynamic Response Function (HRF) (see Methods). The left hemisphere cortical surface (left), and subcortical axial slices (center: thalamic and cerebellar view) show where pulses were detected in each of the participants (Nico, top; Ashley, middle; Omar, bottom). The color scale spans 2 seconds bracketing the average left SM1_ue_ pulse peak. The maps display the top 20^th^ percentile of highest pulse detectability. The participant-specific upper extremity somatomotor region is outlined in black (left). On the right the individual (thin lines) and average (thick line) pulse timecourses are shown (y-axes: percent signal change) for the left SM1_ue_, the left thalamus and right cerebellum.

Disuse pulses propagate through the brain in a specific temporal sequence. We previously reported that SMA regions peak earlier than left SM1_ue_, followed by the cerebellum. On average, the central thalamus pulses peaked later than in left SM1_ue_ (Figure 2; first and second columns, Nico +0.75 seconds (sd 0.16); Ashley +1.07s (sd 0.02); Omar +0.95s (sd 0.09)).

### FC changes and disuse pulses overlap in the central thalamus

Figure 3 illustrates the topography of pulses in relation to FC changes. All participants showed overlap (green) between disuse-driven FC changes and pulses in the dorsal medial cortex (SMA, pre-SMA, dACC), the central thalamus and the effector-specific motor regions of the cerebellum (Figure 3 and Figure S3). Ashley’s pulses in the posterior putamen also overlapped with FC increases.

**Figure 3:**
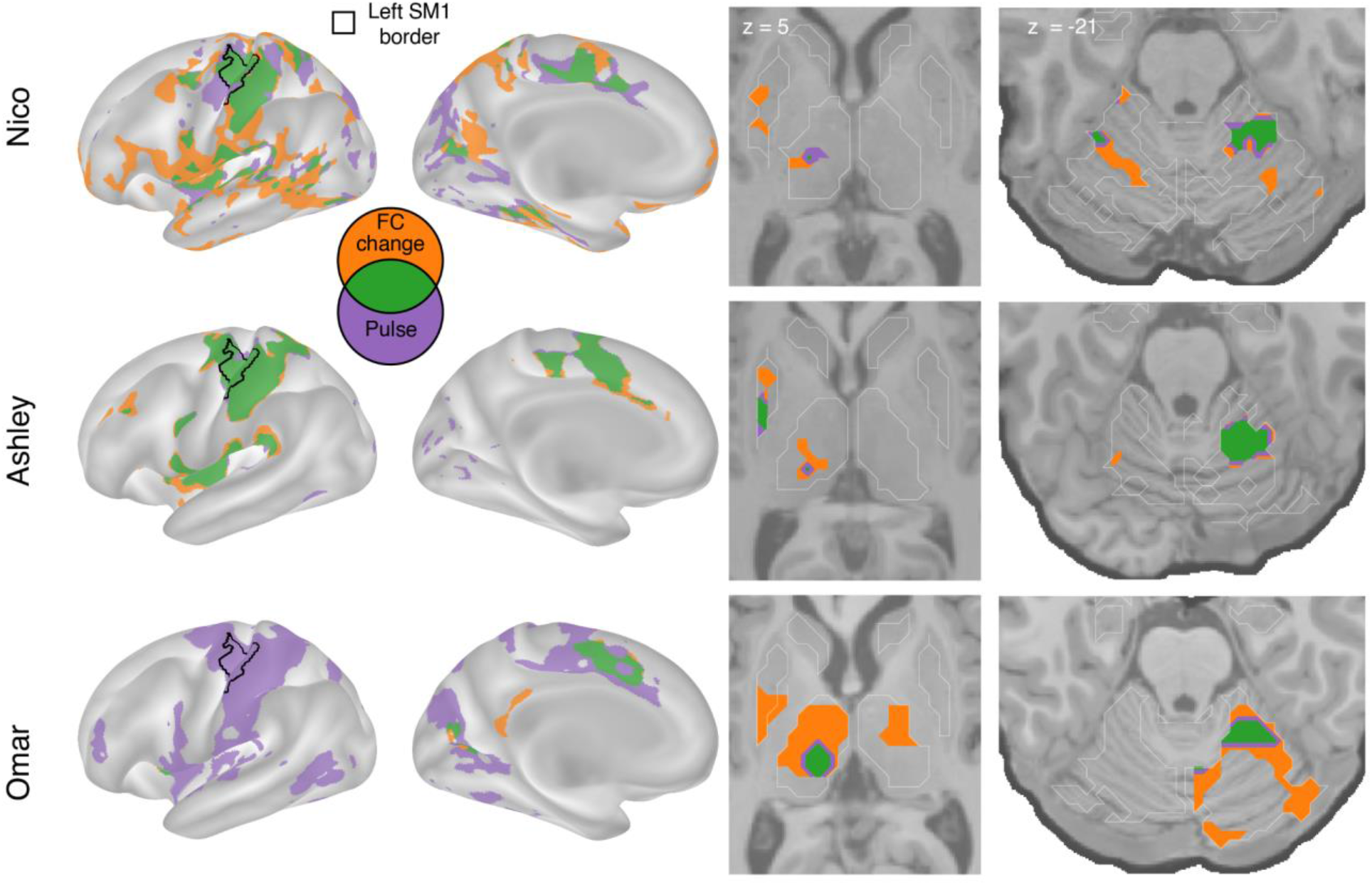
Spatial overlap of functional connectivity (FC) increases and disuse pulses. The strongest disuse-driven FC increases (orange, Cast > Pre, cluster corrected) and disuse pulses (purple, top 20% threshold), as well as their overlap (green) are shown on the cortical surface (left), in the thalamus and putamen (middle) and the cerebellum (right). Results are displayed on the lateral left hemisphere surface, medial left hemisphere surface, and two axial slices (MNI z = 5 and -21). White borders on axial slices defined individual specific FreeSurfer based anatomical structures (z = 5: putamen, globus pallidus, caudate, thalamus; z = -21: cerebellum, hippocampus).

### Subcortical disuse-driven plasticity overlaps with fMRI task activations

To determine whether FC changes and disuse pulses spatially coincided with regions active during upper extremity movement, we analyzed motor task fMRI data collected at baseline (Figures 4,5). The motor task includes simple hand, tongue, and foot movements in a block design (same as used in the Human Connectome Project). This paradigm elicited somatotopic specific responses in the primary motor cortex (see Figures S6-9).

**Figure 4:**
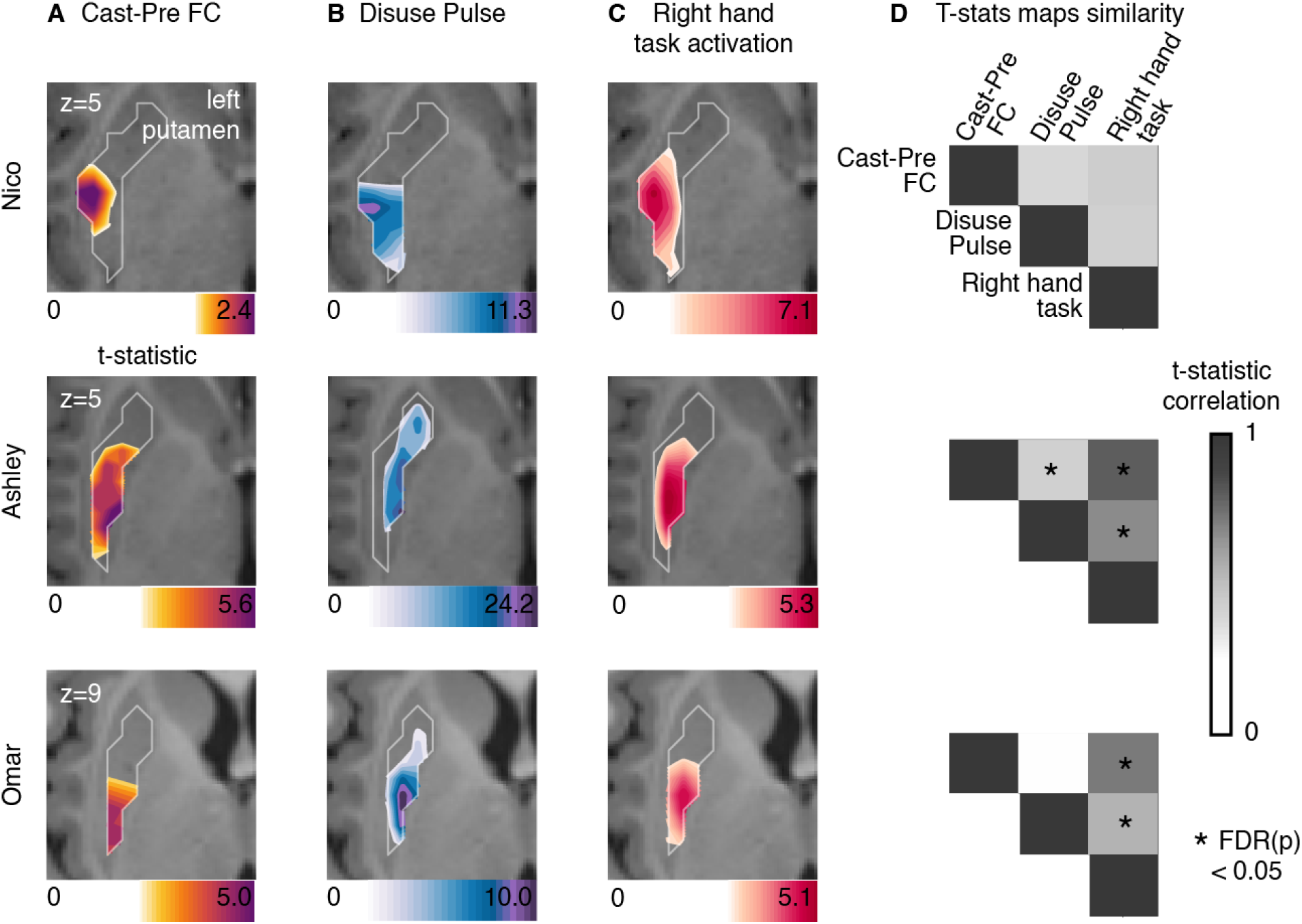
Putamen representations of disuse-driven FC changes, pulses and hand movement task fMRI activations. (A) Map of disuse-driven increases in FC with the left SM1_ue_ region of interest (top 30^th^ percentile t statistics). (B) Map of disuse pulses (top 30^th^ percentile t statistics). (C) Map of pre-casting task fMRI contrast: right hand movement vs baseline (top 30^th^ percentile t statistics). For all three maps, color scales are represented at the bottom of the map with maximum value at 99.5^th^ percentile. (D) Correlation between left putamen t statistic maps for disuse-driven FC increases, pulse and activation during hand movement (right hand vs baseline). Correlations between unthresholded t statistics maps were tested against individual-specific null distribution effects for each participant (top to bottom: Nico, Ashley, Omar). Reported significant p < 0.05 corrected for FDR (black *).

**Figure 5:**
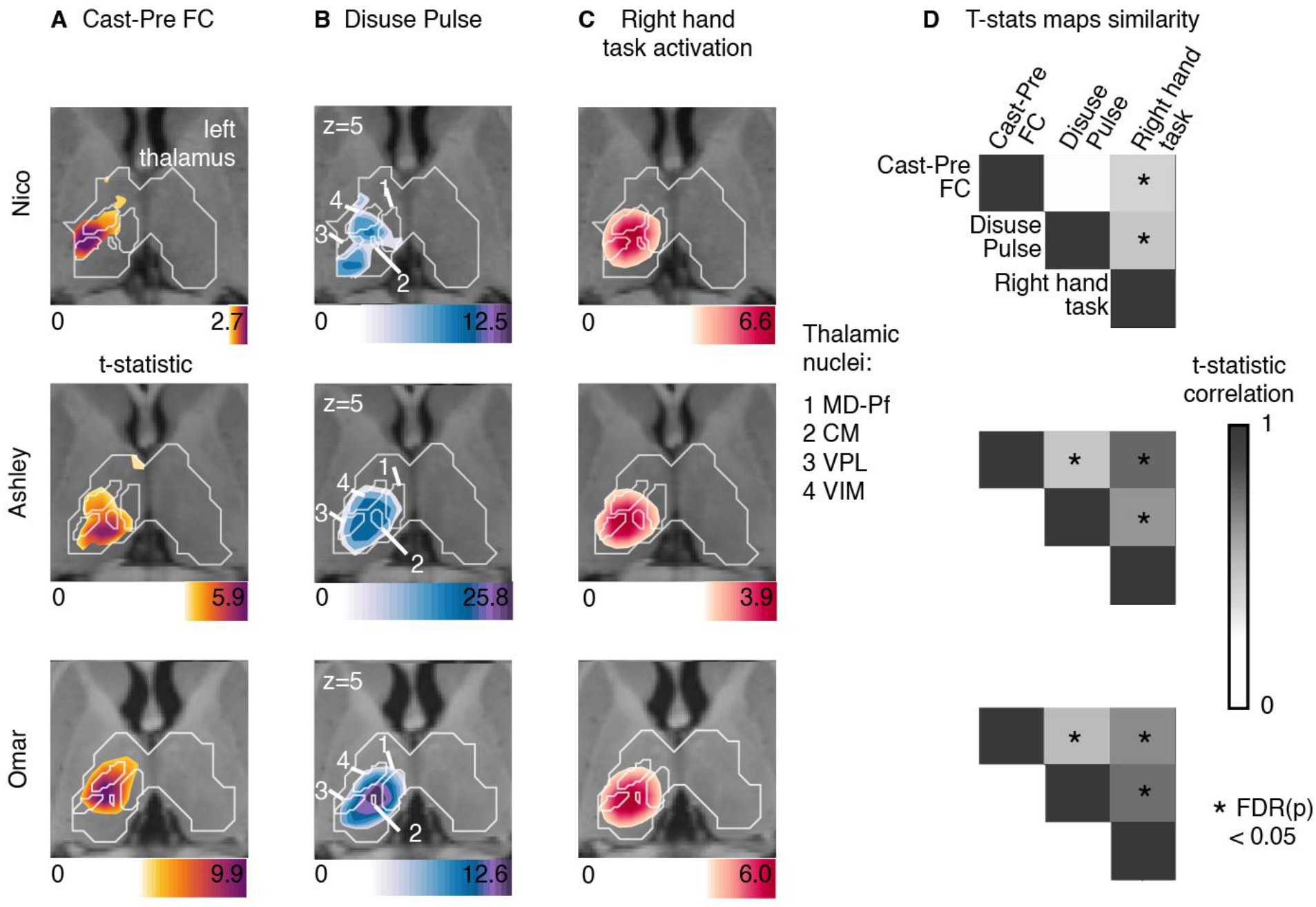
Thalamus representations of disuse-driven FC changes, pulses and hand movement task fMRI activations. (A) Map of disuse-driven increases in FC with left SM1_ue_ region of interest (top 30^th^ percentile t statistics). (B) Map of disuse pulses (top 30^th^ percentile t statistics). (C) Map of precasting task fMRI contrast: right hand movement vs baseline (top 30^th^ percentile t statistics). For all three maps, color scales are represented at the bottom of the map with maximum value at 99.5^th^ percentile. For all three maps, color scales are represented at the bottom of the map with maximum value at 99.5^th^ percentile. Nucleus borders from the THOMAS individual-based segmentation overlapping with the effects are visible in white. Four nuclei are displayed: CM: centro-median, VPL: ventro-posterior lateral, VIM(VLPv): ventral intermediate (ventro-lateral posterior ventral), MD-Pf: medio-dorsal. Average t-statistics values for all thalamic nuclei are represented in bar plots (see figure S4) (D) Correlation between left thalamic t statistic maps for disuse-driven FC increases, pulse and activation during hand movement (right hand vs baseline). Correlations between unthresholded t statistics maps were tested against individual-specific null distribution effects for each participant (top to bottom: Nico, Ashley, Omar). Reported significant p < 0.05 corrected for FDR (black *).

All three participants showed motor task fMRI activations in the posterior portion of the putamen during right hand movement (Figure 4). In each participant, task activations overlapped with the FC change and disuse pulses. This similarity was confirmed by the strength of correlation between t-statistic maps. All participants showed strong (significant for Ashley and Omar) topographic similarities between motor task responses, FC change, and pulse density maps. Statistical significance of map similarity (Figure 4D) was assessed using individual-specific null distributions (see Methods).

All three participants showed task fMRI activations in central thalamus (Figure 5). Task activations overlapped with FC changes and disuse pulses in each participant. This similarity was confirmed by the strength of correlation between t-statistic maps, showing significant similarity in comparison to spatial null distributions (see Methods, Figure 5D). Individual differences in effect sizes were similar in all three measures (as in Table 1). Overall, thalamic FC changes and disuse pulses were similar to task fMRI activations when moving the right hand.

To delineate the specific thalamic distribution of cast-induced plasticity effects in comparison to hand motor circuitry, we quantified the average t-statistic values within each thalamic nucleus from the THOMAS atlas (Figure 5, S4, individual THOMAS atlas segmentation, see Method). Thalamic nuclei showing the greatest average t-statistic for both plasticity effects were the centro-median (CM), the ventroposterior lateral (VPL) and ventral intermediate (VIM) (see Figure S6 for all participants). Right hand movement showed consistent activations of CM, VPL, VIM across all participants. This set of nuclei remained specific when quantifying t-statistic average values from right > left hand movement task contrast, thus confirming the specificity of the right-hand movement circuitry (Figures S7-9). CM, VPL and VIM showed disuse pulse and increased FC changes with left SM1_ue_.

## Discussion

### Disuse driven plasticity effects are not limited to cortex

Using analyses optimized for characterizing fMRI signal differences across the cortex and subcortex, we observed significant subcortical FC changes and disuse pulses in the central thalamus (VIM, CM) and posterior putamen as a result of dominant upper extremity disuse, due to casting. These findings suggest that disuse can drive network changes at all levels of motor circuitry: cortex, cerebellum, thalamus and striatum. In the cortex, the SCAN regions do not show an increase in FC or pulse presence across participants, which highlights the functional differences between SCAN and effector specific networks. Throughout motor and action control circuits, as verified by task fMRI activations, FC increases partially overlapped with the presence of disuse pulses, suggesting that they might represent a brain-wide disuse plasticity mechanism. We previously demonstrated that the presence of disuse pulse does not fully explain the changes in FC^16^. While the presence of disuse pulses in the thalamus suggests a more general phenomenon, other plasticity mechanisms for neuronal and synaptic adaptation likely act in parallel.

### Disuse strengthens FC in motor and action sub-circuitry

Increased FC between subcortical and cortical motor regions, in the presence of pulses, is consistent with our previously reported finding of strengthened FC between motor cortex and the CON ^16^. However, these results cannot be fully accounted for by a simple Hebbian-like process ^39,49,50^, which one would predict should weaken FC in the setting of disuse (not firing together). Instead, we speculate that pulses, by creating co-activation in the disused motor control circuit, could help within network synchronization and information transfer, therefore playing a role in plasticity in order to help maintain the integrity of disused subcircuits ^17^. Pulses in the disused subcircuits may enable the rapid recovery of both behavior (actigraphy) and functional connectivity (within days) after cast removal ^17^.

Alternatively, the circuit can be going through integration of new motor programs related to disuse. Prior work in animal models suggests that neural plasticity in response to changes in auditory or visual stimuli starts with a reduction of inhibitory interneuron activity at the population level, increasing excitatory activity which facilitates Hebbian processes ^37,51–53^. Therefore, the observed FC increase within the motor execution circuit could indicate an increased simultaneous firing, eliciting Hebbian plasticity and information retention. This rise in local activity could increase the likelihood of spontaneous activity pulses, whether they represent runaway activity or necessary homeostatic regulation processes or windows for cortico-thalamic information exchange.

### Disuse-driven plasticity affects posterior putamen involved in habit formation

Rapid, goal-directed learning is thought to primarily involve the dorsomedial striatum, including the caudate, whereas the slower acquisition of habits, which are insensitive to changes in the reward value of the outcome, is thought to depend more on the dorsolateral striatum, including the posterior putamen ^53–56^. The increased FC with the posterior putamen suggests that it might specifically be related to protecting existing motor skills or acquiring new ones. Indeed, the first day removing the cast, our participants kept using their non casted arm more than their recently freed dominant arm, suggesting that participants learned to suppress movements of the casted arm. This trend disappeared on day 2 after cast removal and no fine motor or coordination motor impairment the day of cast removal.^17^

### Plasticity in thalamic nuclei for motor execution

The central thalamus is important for motor and action execution and studies have tied movement performance to activity in VIM, VPL and CM thalamic nuclei ^6,57^, which is consistent with our motor task fMRI activations (Figure S4, S6-8). The VIM is part of the thalamic ventro-lateral region (VL), primarily involved in motor control and relays information between the basal ganglia, cerebellum and motor cortex, whereas the VPL is part of the somatosensory system and relays sensory information related to touch, temperature, and pain from the body to the primary somatosensory cortex ^58–60^. VIM is a common and effective DBS target for tremor. In cases of neurological injury or disease, such as stroke or neurodegenerative disorders, the VIM can undergo plasticity as part of the brain’s adaptation and recovery mechanisms ^36,61^. In rats, for example, VIM stroke results in specific skilled locomotor impairment (e.g., impaired ladder mobility, but not flat walk) ^62^. Reorganization of activity in VIM after the stroke correlates with restored mobility ^63,64^. In fact, successful movement performance, such as walking on a complex surface, after a VIM partial lesion are compensated within a few days, showing the highly effective plasticity of the thalamus ^62^. While the thalamus is essential to cortical organization and specialization during development^65^, plastic reorganization of specific motor programs can also occur in response to motor demands driving functional adaptation in adults. For example, in humans, in the context of amputation, one study observed changes in the representation of the affected body parts within the VIM. Electrophysiological recording in a patient with arm amputation and long-term use of a prosthetic has shown an enlargement of the shoulder representation in the VIM, which now is used for prosthetic grip control by the patient ^36^.

### Thalamic plasticity pulses could carry local sleep spindles

In addition to the VIM and VPL, the CM also showed FC increases, pulses and task fMRI activation. The CM plays a special role in regulating arousal ^66^. While the VIM is the DBS target of choice for treatment of tremor, the CM is targeted in the treatment of Tourette syndrome and, with increasing frequency, also epilepsy and disorders of consciousness ^67^.

Indeed, the thalamus has a specific role in sleep pressure and wakefulness regulation^68^. The thalamus regulates sleep stages and slow oscillations in deep sleep ^45^. Sleep events like thalamo-cortical spindles are thought to help memory and skill consolidation ^69–71^. During deep sleep, slow waves of activity are observed across the cortex and relayed in the thalamus ^72^. These slow waves are thought to help with homeostasis of neural activity after a day of novel experiences ^49,73,74^. The slow waves help networks to synchronize and increase phase locked communication between regions ^75,76^.

The occurrence of disuse pulses in the central thalamus, most reliably in the CM, raises the question whether they might be related to thalamocortical sleep spindles. Could the disuse pulses represent a circuit-specific, sleep-like phenomenon happening during the awake resting-state? Indeed, local sleep can be observed during wakefulness ^77^. After a long period in an awake state, EEG recordings in awake rats can capture local ‘offline’ stages similar to sleep, together with slow waves ^78,79^. Consistent with this idea, the one participant who was always scanned in the morning (Nico) also had the lowest number of pulses. In contrast, Ashley and Omar were always scanned in the evening. Thus, hours spent active since awakening could have driven the frequency of circuit-specific spontaneous activity pulses at the time of scanning.

### Cortico-striato-thalamic FC increase as a marker of disuse in Parkinson’s Disease

In our arm immobilization paradigm, we observed increases in FC in the disused motor circuit, but after removing the cast, these FC changes reversed rapidly and motor behavior was unimpaired^16^. In Parkinson’s disease, action and motor impairments include tremor, rigidity and bradykinesia on top of a general paucity and slowness, and follow subcortical progressive loss of dopamine neurons projecting then affecting the caudate and putamen ^5^. Similarly to our cast experiment, patients with Parkinson’s disease show an increase in FC between putamen, central thalamus and cortical motor area, especially during akinesia ^80–89^. This suggests a potential pathophysiological link between limb disuse and Parkinson’s disease. If increased FC in the motor and action circuits is a marker of increased neuronal excitability when a brain circuit is idling ^90–93^.

VIM is a target for DBS treatment of tremor. Electrophysiological studies in patients with Parkinson disease and parkinsonian mouse models have revealed increased beta power and prolongation of beta burst discharges propagating throughout cortico-basal basal ganglia circuits ^94–97^. These beta bursts disappear when moving or with L-dopa treatment ^98–100^. Further research is needed to determine whether beta bursts detected with electrophysiology are Parkinson’s specific or perhaps disuse-related. Through this parallel, a potential role of disuse pulses would be to behave like sleep thalamic spindle, that are characterized by high frequency such as beta^101^, and to open synchronization windows for information updating in the motor network.

### Plasticity and stability in subcortex

Disuse-driven FC changes and spontaneous activity pulses are not confined to the cortex but extend into the putamen and central thalamus (VIM, CM, VPL). The anatomical pattern, especially the prominence of the CM evokes parallels to sleep-related mechanisms of consolidation and plasticity, as well as to the beta bursts seen in Parkinson’s patients. These findings open up intriguing new avenues for studying disorders such as Parkinson’s disease and sleep physiology. They also raise the interesting possibility that the mechanisms for changing the brain and for maintaining it are one and the same.

## Methods

### Human participants

We analyzed a previously published dataset that comprised three healthy adult volunteers. The first participant (Nico) was 35 years old at the time of scanning and is male. The second participant (Ashley) was 25 years old and female. The third participant (Omar) was 27 years old and male. All participants were right-handed, as assessed by the Edinburgh Handedness Inventory ^102^ (Nico: +100, right handed; Ashley: +91, right-handed; Omar: +60, right-handed). The Washington University School of Medicine Institutional Review Board approved the study protocol and provided experimental oversight. Participants provided informed consent for all aspects of the study and were paid for their participation.

### Experimental setup

Arm immobilization was conducted by constraining the participant’s dominant (right) arm for two weeks (cast period). The immobilization followed a two-week experimental baseline (pre-cast period) and was followed by a recovery period of two weeks (post-cast period). For one participant (Nico), the pre-cast period was acquired one month before the cast period and data was consistently acquired at 5 AM, while for the two other participants, fMRI was acquired at 9 PM. For one participant (Omar), the cast was removed and reapplied after one day of the cast period to adjust for finger comfort. Details of cast construction are described in Newbold et al. ^17^

### Imaging data

On each day of the experiment, a scan session was conducted to acquire structural and functional data. Structural MRI was consisted of four T1-weighted images (sagittal acquisition, 0.8 mm isotropic resolution, 3D MP-RAGE, Gradient echo) and four T2-weigthed images (sagittal acquisition, 0.8 mm isotropic resolution, 3D T2-SPC, Spin echo). A 30 minutes resting state fMRI (rs-fMRI) run was acquired during each session, and two runs of the HCP Motor strip mapping task ^103,104^ were acquired for each pre and post cast session. Ashley and Omar’s fMRI was acquired with an improved fMRI sequence (all 2D Gradient echo, echo planar, TR: 1.1 vs. 2.2 seconds in Nico, 2.6 vs. 4 mm isotropic resolution in Nico). All data were resampled at 3mm in atlas space. Acquisition parameters and procedures are detailed in Newbold et al. ^17^.

### Precision functional analysis

All following data processing and analysis is conducted at the participant level using individually defined functional and anatomical boundaries. All statistical testing was done against null distributions built for each participant. All major results were replicated across all participants.

### MR Image Processing

Preprocessing of structural and functional images, denoising of rs-fMRI data, and cortical surface projection were performed as previously described^17^. Functional image processing followed a previously published pipeline ^105^ and involved temporal interpolation to correct for differences in slice acquisition timing, rigid-body correction of head movements, correction for susceptibility inhomogeneity-related distortions, and alignment to atlas space. The present results are reported in MNI152 space. For cortical surface projection and creation of cifti images, individual-specific surfaces were created defining the cortical gray-matter boundaries derived from T1-weighted images using FreeSurfer ^106^. Subcortical boundaries from the FreeSurfer segmentation were used to select voxels of interest for building the volume part of the cifti image. There were no systematic differences in head movement (mean FD or number of frames removed) between phases of the casting protocol.

rs-fMRI data denoising involved replacement of high-motion frames (framewise displacement [FD] > 0.1 mm) by temporal linear interpolation, band-pass filtering (0.005 to 0.1 Hz), and regression of nuisance time series, including head movement parameters, the global signal averaged across all gray-matter voxels, and orthogonalized waveforms extracted from ventricles, white matter, and extracranial tissues. The rs-fMRI for functional connectivity is then projected to cortical surface as the last step and smoothed using a two-dimensional 6-mm full-width half-maximum (FWHM) smoothing kernel and volume data were smoothed using a three-dimensional 4.7-mm FWHM kernel.

Hemodynamic response function modeling was used for the pulse spatio-temporal representation analyses using rs-fMRI and for the HCP task data. For these analyses, denoising of data involved high-pass filtering (0.1Hz) before surface projection and regression of nuisances time series (head movements parameters, individual high-motion frames). Nuisance regression was done at the surface and subcortical volume level within a GLM design. Cortical surface data were smoothed at the last stage of each analysis using a two-dimensional 6-mm full-width half-maximum (FWHM) smoothing kernel and volume data were smoothed using a three-dimensional 4.7-mm FWHM kernel.

Fully processed data are available in the Derivatives folder of the Cast-Induced Plasticity dataset in OpenNeuro (https://openneuro.org/datasets/ds002766).

### Primary somatomotor upper extremity region of interest

The upper extremity SM1 region was defined in individual using a task-based approach combined with automatic labeling by FreeSurfer. Details were previously described in Newbold et al. ^17^.

### Individual network representation

A set of 18 canonical functional networks was defined for each participant using a graph theory-based community detection algorithm with anatomical priors ^20^(see figure S1), Infomap algorithm ^107^(https://www.mapequation.org/). This algorithm assigns grayordinates to communities. Subsequently, these communities are categorized based on their similarity to established group-average networks recently updated to include the SCAN ^20,23^. Final cortical resting state networks were derived from the consensus network assignments obtained through aggregation across thresholds.

### Thalamic nuclei segmentation using THOMAS

The Thalamus-Optimized Multi-Atlas Segmentation (THOMAS v 2.1) method, a promising approach for identifying nuclei, was employed ^28^. The choice of THOMAS followed the latest consensus of nuclei naming and automatic segmentation algorithm improvement. We also aimed to interpret our results in relation to clinical application and needed a robust localization of VIM, which has been shown to co-localized with the segment labeled VLPv (Ventro-Lateral-Posterior ventral) using the THOMAS segmentation ^31^. For the thalamic nuclei segmentation in our precision mapping participant, the hips_thomas.csh function from version 2.1 was utilized. This version has been validated exclusively for T1 acquisition ^108–110^ and is accessible through Docker (https://github.com/thalamicseg/thomas_new). The average T1 acquisition, generated for the registration of all functional data, was employed for this purpose. Nuclei were mapped into cifti format and the resolution of functional image (3mm) to quantify overlap between brain maps and thalamic nuclei.

### rs-fMRI Functional connectivity (FC)

Average BOLD time series were calculated for each individual specific vertices/voxels (i.e., cifti grayordinates) to construct a full brain FC for each rs-fMRI session. Seed FC was estimated as the average of FC maps for each voxel in the seed. Pre-cast FC maps were averaged over sessions to define the baseline FC.

### rs-fMRI FC change

For casting sessions comparison to baseline (pre-cast) sessions, Cohen’s d was calculated at each grayordinate. Cohen’s d indexes change in FC effect size as it accounts for standard deviation. This allows higher sensitivity to subcortical casting effects as the subcortical correlation values are lower in comparison to cerebral cortex but consistent across sessions. Cohen’s d is also more stable than the t-statistic to variation in the number of measurements between participants or between casting protocol phases. Due to repeated measure design, the sample size is small (only 14 sessions per phase). Thus, loss of a single session impacts the interpretation of t-statistic more than Cohen’s d. Statistical significance was assessed within participants using individually generated nulls. To define significant effect, we computed null distribution FC maps by randomizing session labels 1000 times and computing a multi threshold cluster based correction. 10 thresholds were defined within the range of values of the original data, using values of every 10^th^ percentile of the values distribution. Cluster of data passing cluster size correction at p<0.05 for at least two thresholds are displayed in figure 1. Cluster size correction was estimated for each anatomical structure (cortical and subcortical) independently.

### Pulse detection and modeling

To detect large amplitude fluctuation in Left SM1_ue_ regions characteristic to a “pulse”, we look for variation of fMRI activity in Left SM1_ue_ above a standard activity and unilateral. To do so, the average BOLD time series for the Left and Right SM1_ue_ were computed for each rs-fMRI run and normalized across all runs. The difference between Left and Right time series were computed and the 2.3 times the standard deviation of this difference was used as threshold for pulse detection. A Left SM1_ue_ pulse was defined by two criteria. The first criterion was an increase of Left to Right time series difference (above 2.3 times the Left to Right time series difference standard deviation). The second criterion was an increase in signal of Left SM1_ue_ time series above 2.3 times its average time series standard deviation. To avoid movement related to false positives, Pulses with a high correlation (>0.8) with head movement (on a 18s-window centered on the pulse peak) were removed.

In order to study the subcortical pattern of the disuse pulse described in Newbold et al. ^17^, we used a pulse detection analysis sensitive to potential differences in hemodynamic response function (HRF) shape ^111^ in subcortical regions. We modeled, at each grayordinate, a pulse waveform using the HRF shape at each pulse (double gamma HRF function available in nipy (SPM based hrf double gamma, nipy.modalities.fmri.hrf) and optimized using scipy (scipy.optimize.curve_fit)). For grayordinates that did not show a pulse activity, the model did not converge within the parameter range. Regions with the most frequent pulse activity (20% highest pulse detection) are displayed in figure 2. Pulse latency at each pulse locus was computed as the temporal difference between peaks relative to Left SM1_ue_. The displayed map was thresholded at 1.1 seconds (TR) after the Left SM1_ue_ pulse peak.

### Motor Task analysis

Task block designs for each movement condition (Tongue, Left Hand, Right Hand, Left Foot, Right Foot) were modeled using a double gamma HRF in a GLM analysis from FSL feat^103^. Block onset and offset were modeled as independent events ^112^. Analysis was conducted independently on surface and subcortical grayordinates, following the HCP pipeline steps (release v4.3). Second level analysis across runs within participants was performed with FSl and the resulting t-scores were used to study thalamic responses.

### VIM localization

Voxels representing VIM were determined using the anterior commissure - posterior commissure (ACPC) formula ^113^ as follows. The T1 was aligned on the ACPC line using ACPC detect ^114^. We used the FreeSurfer segmentation of the third ventricle to estimate VIM coordinates. The coronal range was estimated as 1/4 of the third ventricle length from the posterior limit of the ventricle with 2 mm anterior range. The sagittal range was estimated between 14 mm from the center of the third ventricle and 11 mm from the border of the third ventricle. The voxels between the three axis range were brought to functional orientation and to the functional resolution of 3mm.

### Quantification and testing of effects against individual null distribution

To test the significance of results per anatomical region (e.g. subcortical structure or thalamic nuclei), given spatial autocorrelation in BOLD signal, we used null distribution testing. We generated 1000 random representations at the grayordinates level, using Moran spectral randomization, an algorithm that gets informed by spatial distances between vertices ^115,116^. We compute the values of interest, i.e. average values per nuclei or correlation across voxels of the thalamus and compare the true to the random null distribution data. Tests were corrected for multiple comparisons across participants using false discovery rate.

## Acknowledgments

This work was supported by NIH grants NS123345 (B.P.K.), NS098482 (B.P.K.),MH129616 (T.O.L.), T32DA007261 (S.R.K), MH096773 N.U.F.D.), MH122066 (N.U.F.D.), MH121276 (N.U.F.D.), MH124567 (N.U.F.D.), NS129521 (N.U.F.D.), and NS088590 (N.U.F.D.); by the Taylor Family Foundation (T.O.L.); by the Intellectual and Developmental Disabilities Research Center (N.U.F.D.); by the Kiwanis Foundation (N.U.F.D.); by the Washington University Hope Center for Neurological Disorders (B.P.K. and N.U.F.D.); and by Mallinckrodt Institute of Radiology pilot funding (N.U.F.D.). Computations were performed using the facilities of the Washington University Research Computing and Informatics Facility, which were partially funded by NIH grants S10OD025200, 1S10RR022984-01A1 and 1S10OD018091-01. Additional support is provided by the McDonnell Center for Systems Neuroscience.

