## Supplementary figures for "Disuse-driven plasticity in the human thalamus and putamen"

### Supplementary Figure

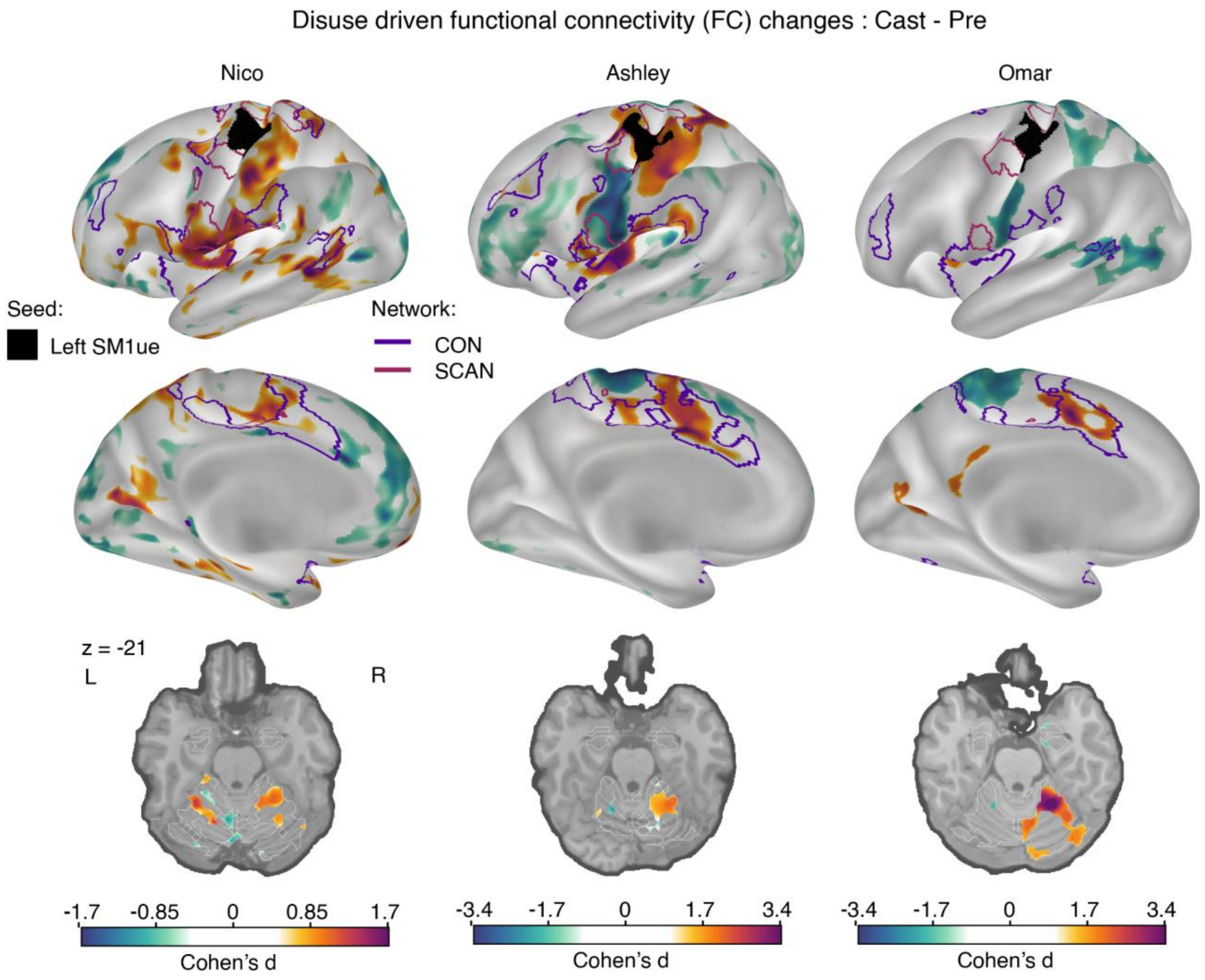


***Figure S1: Disuse-driven changes in functional connectivity (FC) of effector-specific primary somatomotor cortex (L-SM1$_{ue}$) in cortex and cerebellum.*** *Individual-specific plasticity effect size (Cohen's d) maps showing changes in FC during casting (Cast − Pre) for the L-SM1$_{ue}$ (black), and for each participant (left to right columns: Nico, Ashley, Omar). For reference, a Cohen's d of 0.8 is generally considered a large effect size. Only significant effects after cluster correction at $p < 0.05$ (see Methods) are displayed. With a TR that is twice as long, Nico's effect sizes are about half the size of the other participants. The functional network borders of the Cingulo-opercular (CON, purple) and Somato-cognitive action (SCAN, maroon) networks are displayed on the inflated surface rendering (top row). The freesurfer based anatomical border of cerebellum is overlaid on the axial slices(bottom row).*

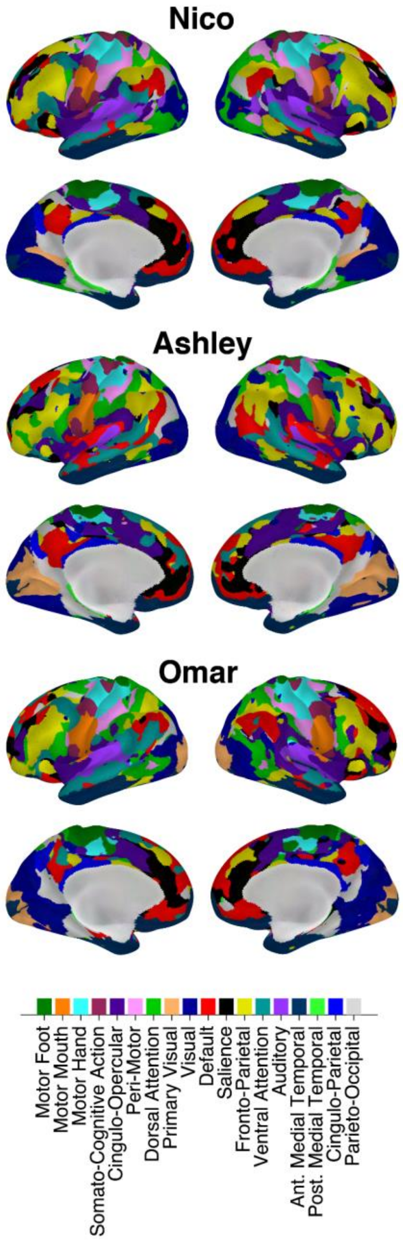


***Figure S2 individual infomap in the cortex.*** *Infomap-based individual definition of the 18 canonical networks for each individual (Nico, top; Ashley, middle; Omar, bottom)*

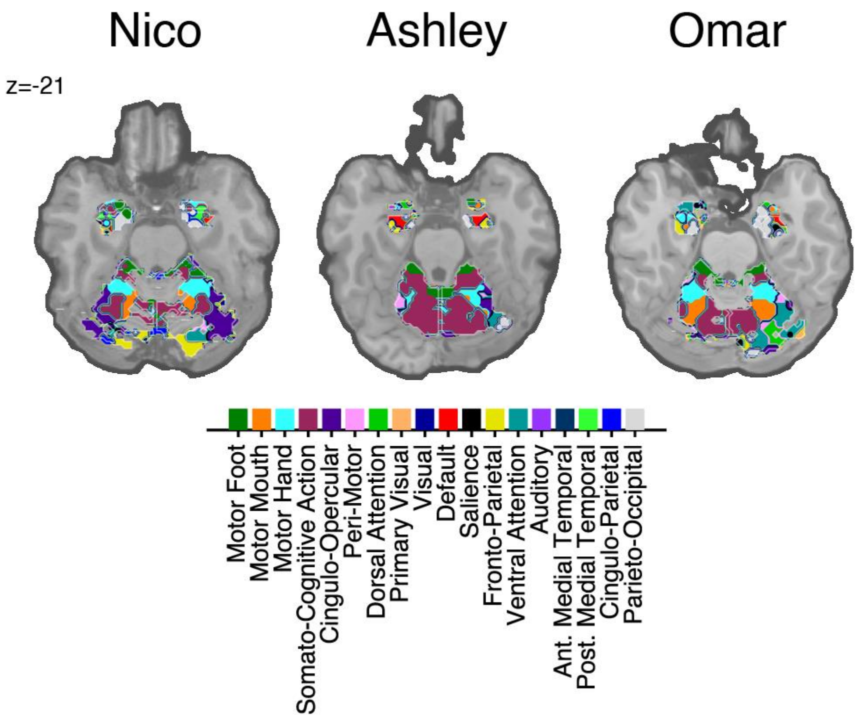


***Figure S3 individual infomap in the cerebellum.*** *Infomap-based individual definition of the 18 canonical networks for each individual (Nico, top; Ashley, middle; Omar, bottom)*

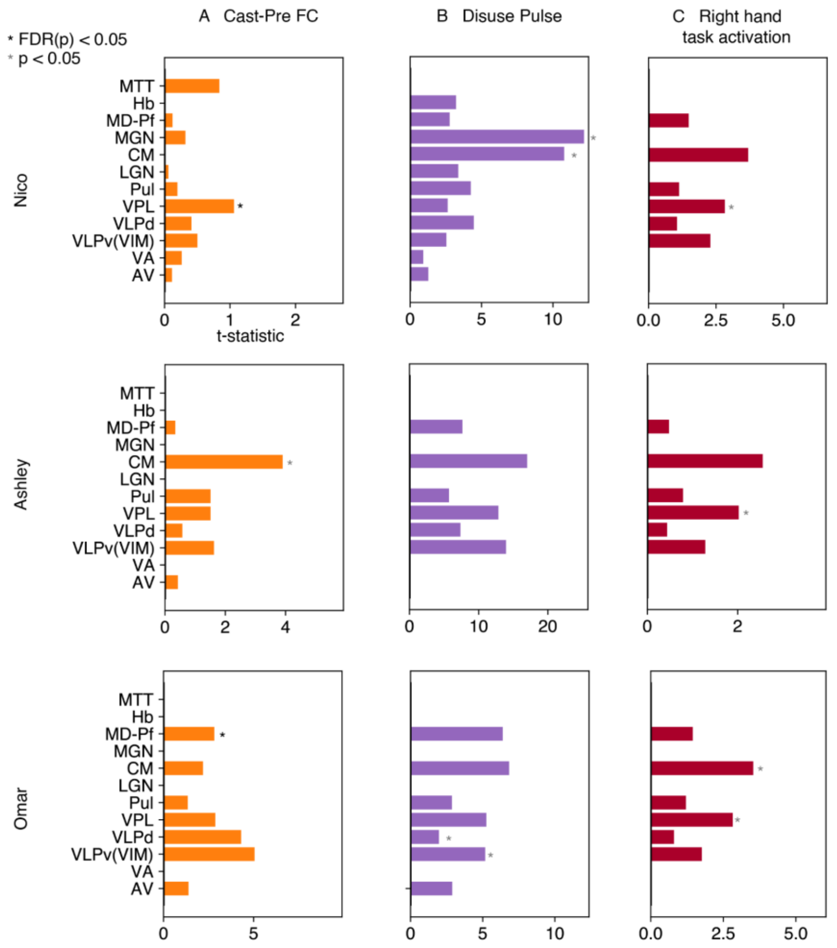


***Figure S4: Average t statistics per left thalamic nuclei of the THOMAS atlas individual segmentation*** *(A) top 30th percentile t statistics left thalamic map of increase FC during casting with Left $SM1_{ue}$ region (B) top 30th percentile t statistics left thalamic map of disuse pulses during casting (C) top 30th percentile t statistics left thalamic map of Right hand movement vs baseline contrast of the HCP motor task from pre casting performance. Significance testing is performed using effect specific null distribution (see Method). * indicates p values <0.05, in black if passing false discovery rate, in gray if not. (CM: centro-median , VPL: ventro-posterior lateral , Hb: habenula, VLPv(VIM): ventro-lateral posterior ventral (ventral intermediate), MD-Pf: medio-dorsal, Pul: pulvinar, MGN: medial geniculate nucleus, VA: ventral anterior , MTT: mammillothalamic tract, AV: anteroventral, LGN: lateral geniculate nucleus)*

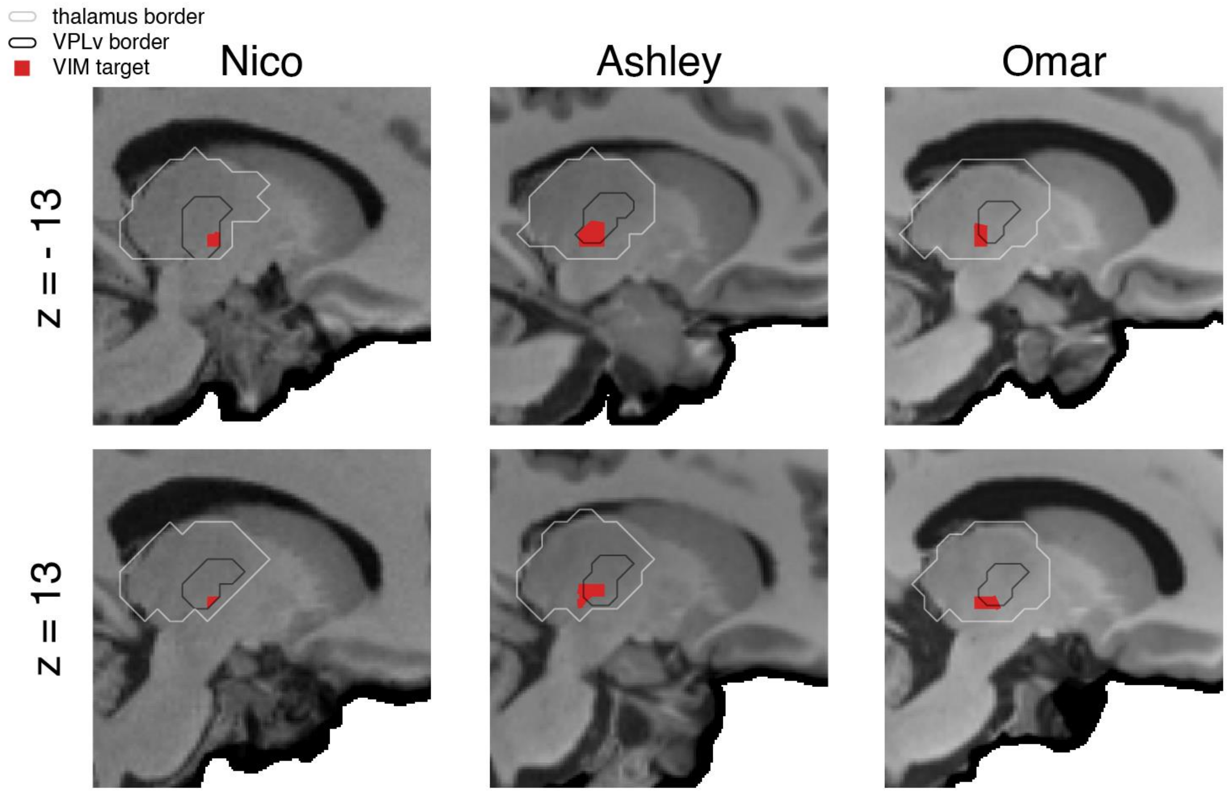


***Figure S5 Localization of Ventrointermediate (VIM) target.*** *The union of possible VIM target voxels (red) within the 2 mm anterior and 3 mm superior of the ACPC target formula. For each participants (Nico, right; Ashley, center; Omar, left), and both hemisphere (Left, top; Right, bottom), the THOMAS segmentation border of the Ventro-Posterior Lateral ventral nuclei (VPLv; black) and thalamic (white) borders are overlaid to highlight the overlap of VIM (target to descend stimulation electrode) with the bottom of the VPLv.*

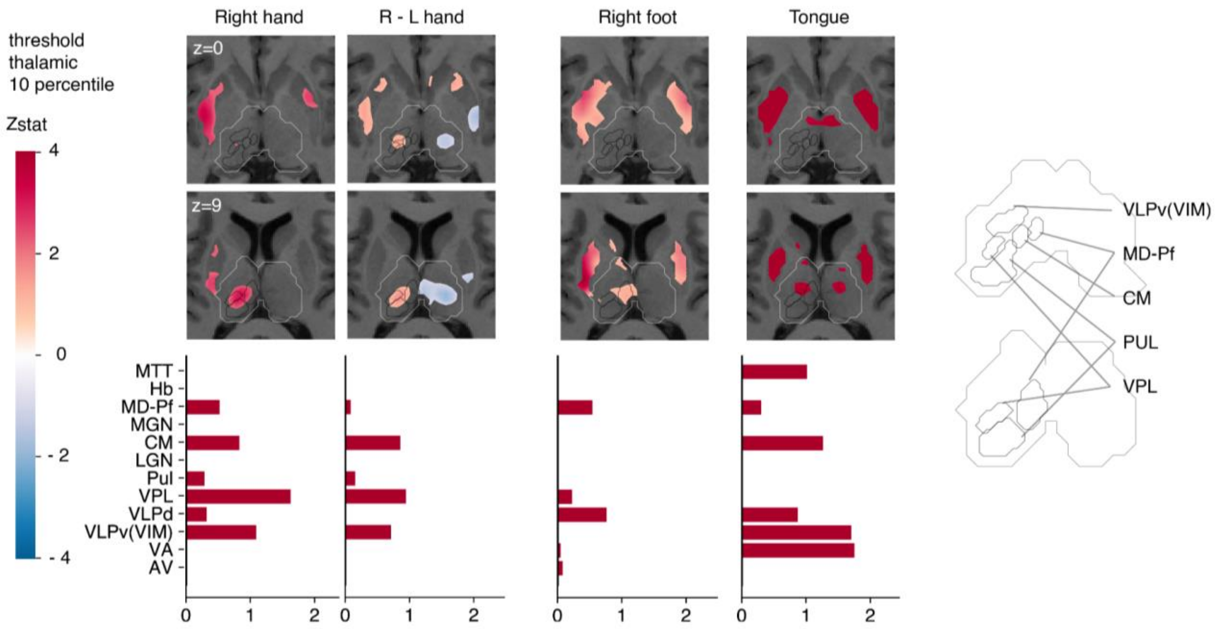


***Figure S6 Subcortical representation of motor task in Nico.*** *Activation maps of HCP motor task contrast, showing the top 10 percentile relative to left thalamic activation (top) and their corresponding average per left thalamic nuclei (bottom). From left to right columns represent contrast of: Right hand vs baseline, Right - left hand, Right foot, Tongue (CM: centro-median , VPL: ventro-posterior lateral , Hb: habenula, VLPv(VIM): ventro-lateral posterior ventral (ventral intermediate), MD-Pf: medio-dorsal, Pul: pulvinar, MGN: medial geniculate nucleus, VA: ventral anterior , MTT: mammillothalamic tract, AV: anteroventral, LGN: lateral geniculate nucleus)*

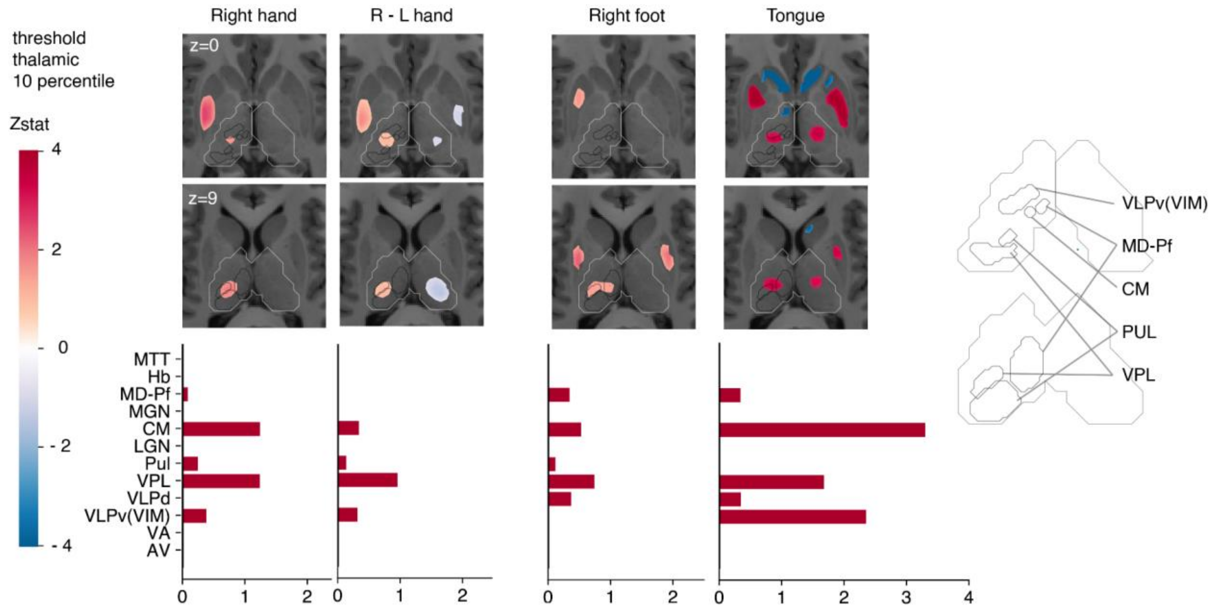


***Figure S7 Subcortical representation of motor task in Ashley.*** *Activation maps of HCP motor task contrast, showing the top 10 percentile relative to left thalamic activation (top) and their corresponding average per left thalamic nuclei (bottom). From left to right columns represent contrast of: Right hand vs baseline, Right - left hand, Right foot, Tongue (CM: centro-median , VPL: ventro-posterior lateral , Hb: habenula, VLPv(VIM): ventro-lateral posterior ventral (ventral intermediate), MD-Pf: medio-dorsal, Pul: pulvinar, MGN: medial geniculate nucleus, VA: ventral anterior , MTT: mammillothalamic tract, AV: anteroventral, LGN: lateral geniculate nucleus)*

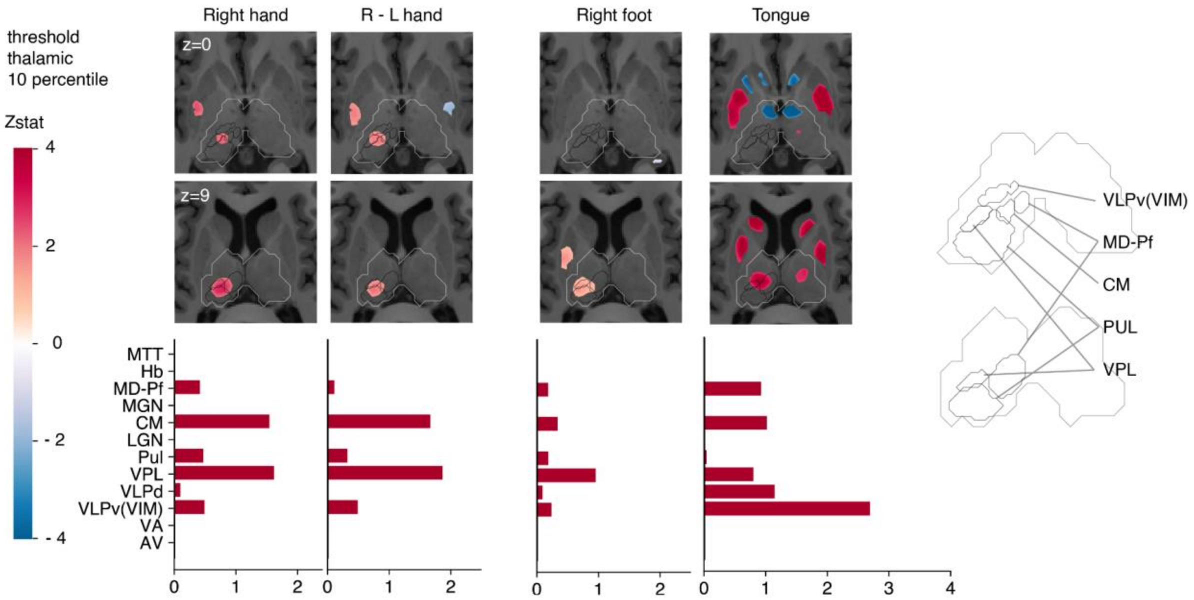

***Figure S8 Subcortical representation of motor task in Omar.*** *Activation maps of HCP motor task contrast, showing the top 10 percentile relative to left thalamic activation (top) and their corresponding average per left thalamic nuclei (bottom). From left to right columns represent contrast of: Right hand vs baseline, Right - left hand, Right foot, Tongue (CM: centro-median , VPL: ventro-posterior lateral , Hb: habenula, VLPv(VIM): ventro-lateral posterior ventral (ventral intermediate), MD-Pf: medio-dorsal, Pul: pulvinar, MGN: medial geniculate nucleus, VA: ventral anterior , MTT: mammillothalamic tract, AV: anteroventral, LGN: lateral geniculate nucleus)*

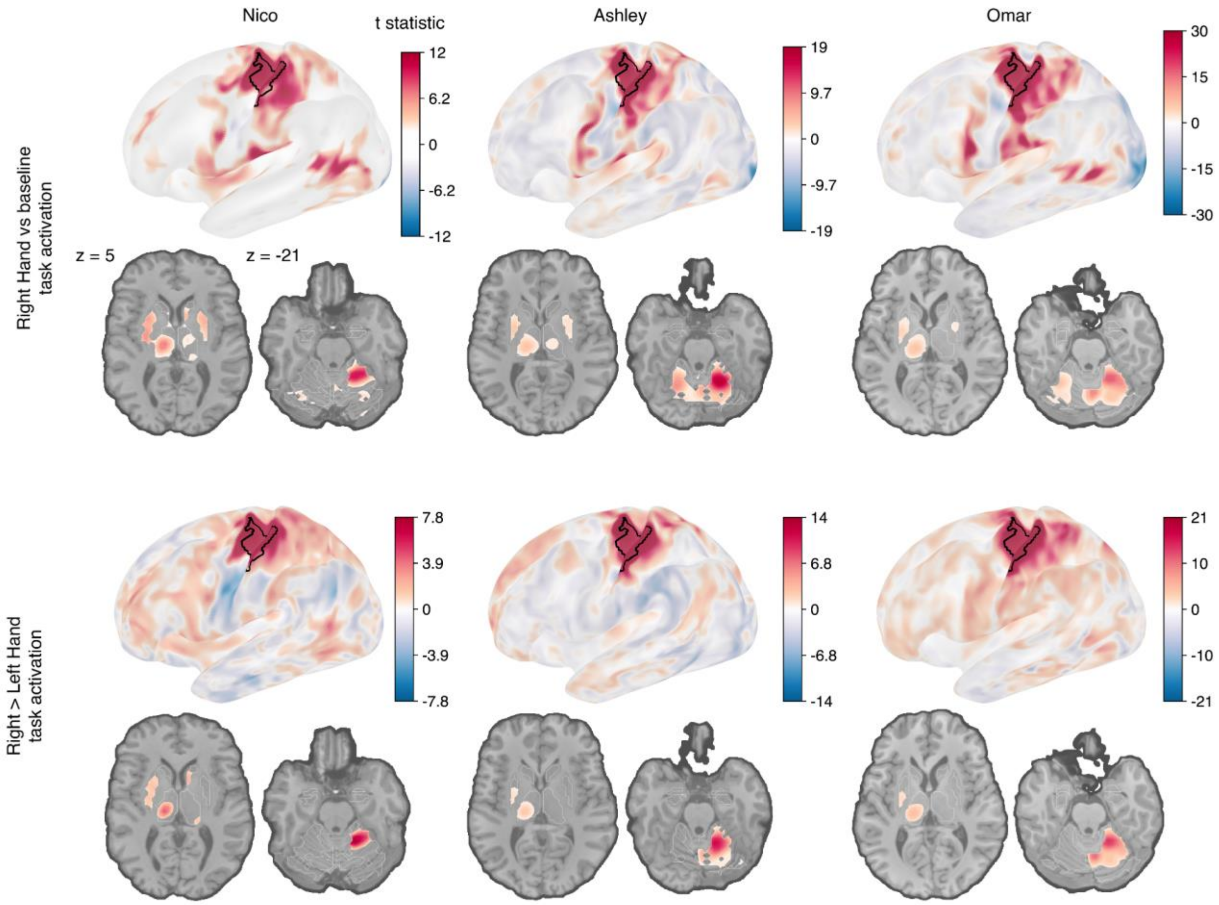


***Figure S9 top 30th percentile of hand motor tasks.*** *T-statistic activation maps of HCP motor tasks contrast Right hand vs baseline (top line) and Right hand > Left hand (bottom line) for each participant. The black border indicates the Left Primary somatomotor area for reference on the inflated surface. axial slices z=% and z=-21 highlights main activation in putamen, thalamus and in cerebellum respectively.*
